# A highly penetrant *LMNA* R541C variant associated with dilated cardiomyopathy leads to dysregulation in metabolism and proliferation pathways in stem cell-derived cardiomyocytes

**DOI:** 10.64898/2026.07.20.739542

**Authors:** Thomas E. Keller, Ci Koehring, Brett R. Higgins, Jiajia Yang, Faiza A. Siddiqui, Forouzandeh Farsaei, Kami Kim, Thomas V. McDonald

**Affiliations:** Division of Infectious Disease and International Medicine, Department of Internal Medicine, University of South Florida; USF Health Heart Institute, University of South Florida; Division of Cardiovascular Diseases, Department of Internal Medicine, University of South Florida

**Author notes:** Corresponding Author: Thomas V. McDonald.

**Keywords:** *LMNA*, cardiomyocytes, cardiomyopathy, RNA-seq

## Abstract

**Background:** *LMNA* codes a widely expressed nuclear cytoskeletal protein (lamin A/C) with multiple important functions. Pathogenic *LMNA* genetic variation may lead to autosomal dominant cardiomyopathy, though the severity and rate of progression can vary with the specific nucleotide change and location. Prior studies showed that induced pluripotent stem cells (iPSC)-derived cardiomyocytes (iCMs) with *LMNA* R541C exhibited reduced *LMNA* protein abundance, increased sarcomere disorganization, and abnormal electrophysiology.

**Methods:** We investigated the *LMNA-*R541C variant that exhibits a highly penetrant and severe clinical cardiomyopathy phenotype using transcriptomic analysis of iCMs. Patient-derived iPSCs with CRISPR-corrected (clustered regularly interspersed short palindromic repeats) isogenic control cells and CRISPR knock-in *LMNA-*R541C heterozygous iPSCs were generated for isogenic controlled experiments.

**Results:** In differential gene expression analyses we observed that *LMNA*^R541C/WT^ iPSC-derived cardiomyocytes had consistent perturbations in 123 genes across CRISPR-corrected and knock-in experiments compared to controls. Pathway analysis identified that the G2M checkpoint and oxidative phosphorylation processes were consistently dysregulated and confirm these findings in previously published iPSC and murine models.

**Discussion:** These results implicate perturbed gene expression and pathways that may contribute to the severe phenotypes in *LMNA-*R541C. Informatic analysis of pathways suggests several drug classes including multiple cardiac glycosides as potential targeted therapeutic candidates to be explored.

## Introduction

Mendelian genetic cardiomyopathies comprise a significant portion of familial heart disease (Hershberger et al., 2013; Peters et al., 2019). Hereditary cardiomyopathies often progress slowly and may remain undetected until the disease has advanced to a severe stage. With increasing recognition and genetic screening of suspect families, more pre-clinical identification of carriers of pathogenic cardiomyopathy variants occurs. Standard heart failure therapies for affected patients may not be appropriate in pre-symptomatic individuals since it is unclear they will impact the earliest molecular and cellular processes triggered by *LMNA* variants. To identify individuals who might benefit from Precision Medicine, early biological pathways need better delineation to distinguish biomarker predictors and molecular mechanisms of disease progression for disease pre-emption.

Pathogenic variants in the *LMNA* gene are often linked to dilated and arrhythmogenic cardiomyopathy (DCM and ACM). *LMNA* encodes the lamin A/C protein that is widely expressed and is a major structural component of the nuclear membrane. It plays important roles in chromatin organization and regulation, transcription regulation, DNA repair, and nuclear-cytoskeletal mechano-transduction (Capell & Collins, 2006; Liu & Ikegami, 2020). *LMNA*-associated DCM is highly penetrant and progressive, where the end-stage frequently involves heart failure and symptomatic cardiac rhythm disturbances (Captur, Arbustini, Bonne, et al., 2018; Crasto et al., 2020). Families and individuals with different *LMNA* mutations often vary in their clinical presentation and rate of progression (Kumar et al., 2016; Peretto et al., 2019). Complete absence of *LMNA* induces severe cardiac dysfunction with early mortality, typically within 8-10 weeks (Auguste et al., 2018; Nikolova et al., 2004). *LMNA*-associated human disease, however, almost universally results from heterozygous (autosomal dominant) inheritance. Pathogenic variants that may be missense or structural (truncations, aberrant mRNA splicing, or indels). Missense *LMNA* variants express mutant *LMNA* proteins that may have pleiotropic aberrant functional properties leading to variable clinical phenotypes and degrees of severity (Captur, Arbustini, Syrris, et al., 2018; Pasotti et al., 2008; van Rijsingen et al., 2012).

We previously described a variety of phenotypic differences arising from specific *LMNA* pathogenic variants using patient-specific induced pluripotent stem cell (iPSC)-derived cardiomyocytes and cardiac fibroblasts (Yang, Argenziano, et al., 2021). Here we have focused on the missense *LMNA*-R541C due to its remarkably severe and early penetrance of heart disease. We identified *LMNA*-R541C in a large family that exhibited aggressively progressive heart failure in multiple adolescent individuals. A separate study has identified two individuals with DCM that carried the *LMNA* R541C mutation and found using a homozygous mouse model that *Lmna*-R541C is associated with ventricular dilation and impaired systolic function(Yang et al., 2022). The early onset of severe heart disease in *LMNA* R4541C highlights the importance of characterizing early regulatory pathways that that may be variant-specific. To date there have been few molecular studies of laminopathies using heterozygous (autosomal dominant) models with mice or iPSCs. Here we study patient-derived *LMNA* R541C iPSC cardiomyocytes using CRISPR-corrected patient-derived iPSCs and CRISPR-based creation of heterozygous *LMNA*-R541C in an isogenic cell line

## Methods

### Pluripotent stem cell culture

The induced pluripotent stem cell (iPSC) line harboring a heterozygous *LMNA*-R541C variant was generated from peripheral blood mononuclear cells (PBMCs) of a patient with severe, familial *LMNA* cardiomyopathy as described (Yang, Argenziano, et al., 2021; Yang, Burgos Angulo, et al., 2021). The subject was recruited under the approval of the Institutional Review Board of the University of South Florida (IRB: Pro00033948). Unrelated control iPSCs without *LMNA* mutation were purchased from ATCC company(Yang, Burgos Angulo, et al., 2021). The control PGP1 iPSCs are a well characterized cell line purchased from Synthego (Synthego Corporation, Redwood City, CA). PGP1 IPSCs were derived from skin fibroblasts from a health Caucasian male (Personal Genome Project GM23248) (Ng et al., 2021).

Cells were cultured in an incubator with humidified atmosphere (5% CO_2_ and 95% air) at 37°C on Matrigel (Corning)-coated plasticware. iPSC growth media was mTeSR-Plus (STEMCELL Technologies). When cell confluency reached 70-80% they were passaged by detaching clumps of cells with phosphate-buffered saline (PBS) with 0.5mM EDTA and replating in mTeSR-Plus supplemented with ROCK inhibitor Y27632(Sigma-Aldrich) overnight.

### iPSC gene editing

To create iso-genic controls, two approaches were used. First, the patient-derived *LMNA*-R541C iPSC line was corrected to wild-type with Prime-Editing (Anzalone et al., 2019; Kim et al., 2021). A complementary approach was used for additional isogenic control where the *LMNA*-R541C variant was “knocked-in” to the control PGP1 iPSC line using CRISPR-Cas9 editing.

For Prime-editing, we modified the pCMV-PE2-P2A-GFP plasmid (Addgene# 132776, (Anzalone et al., 2019)) by replacing the CMV promoter with an EF-α promoter and the P2A-blasticidin resistance cassette from pLenti-PE2-BSD (Addgene# 161514, (Kim et al., 2021))with NEBuilder HiFi DNA Assembly (New England Biolabs) to enhance expression in iPSCs. The resulting plasmid, pEF1α-PE2-P2A-GFP-BSD was confirmed by Sanger sequencing.

Design of the plasmid coding the pegRNA sequences to target and correct the *LMNA*-R541C (c.1621C>T) were determined using Pegfiner (pegfinder.sidichenlab.org, (Chow et al., 2021)). The targeting *LMNA*-R541C correction pegRNA plasmid was designed with the following oligonucleotides:

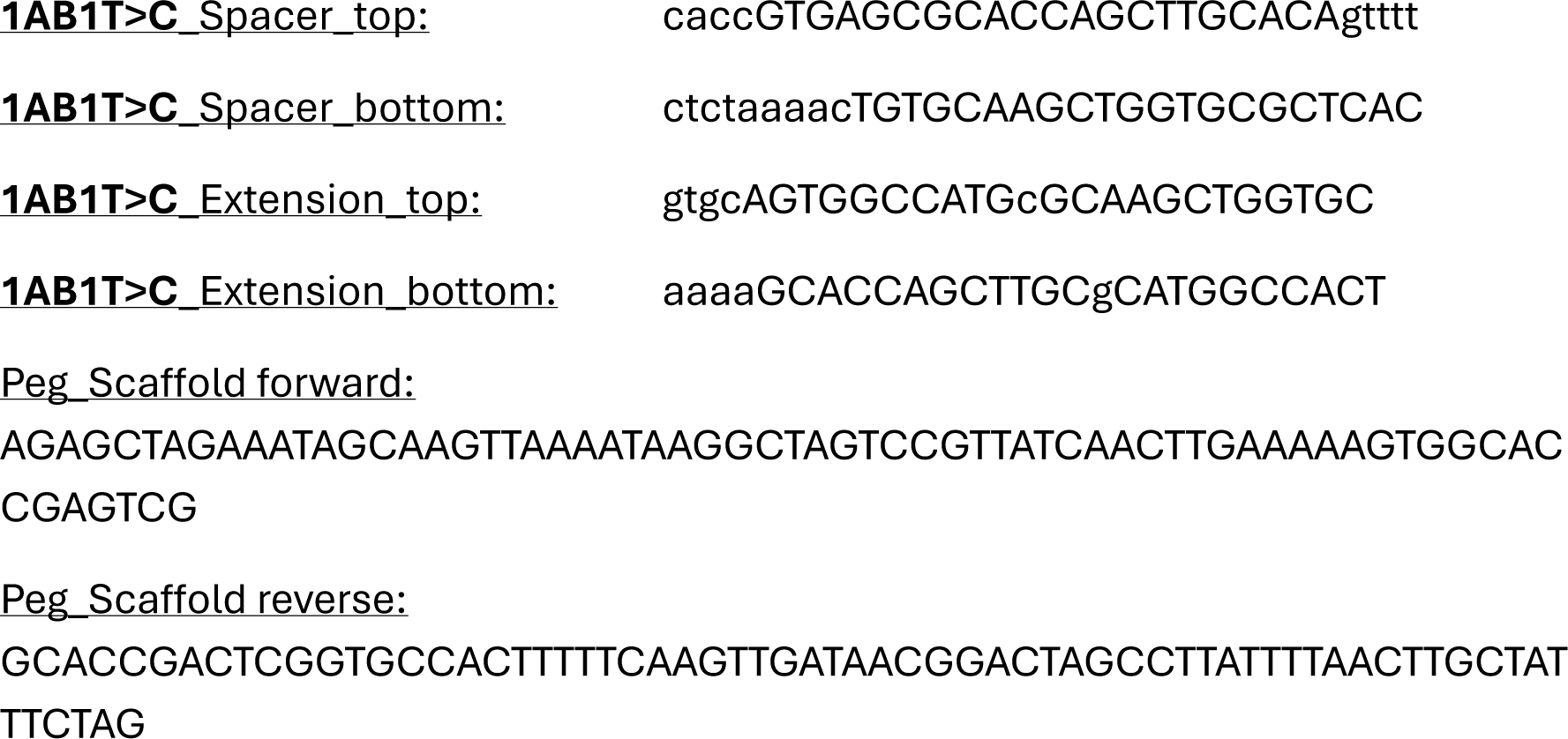

Oligonucleotides were assembled into the pU6-pegRNA-GG-aacceptor plasmid (Addgene#132777) with GoldenGate Assembly (New England Biolabs) to create the plasmid 1AB1-pegRNA and was verified by sequencing. A second plasmid (1AB1-PE3b) that targets adjacent nicking of the non-edited genomic DNA strand for increased editing efficiency was created by ligating the following oligonucleotides into the BsmBI site of plasmid BPK1520 (Addgene#65777, (Kleinstiver et al., 2015)): top-caccGGAAGTGGCCATGcGCAAGC, bottom-aaacGCTTGCgCATGGCCACTTCC.

To achieve Prime Editing, iPSCs growing at log phase were suspended with PBS-EDTA and combined with the following plasmids, pEF1α-PE2-P2A-GFP-BSD, 1AB1-pegRNA, and p1AB1-PE3b (at a ratio of 1.12μg, 0.75μg, and 0.25μg, respectively with1×10^6^cells) in the P3 Primary Cell 4D kit (Lonza). Transfection was done in a Lonza Nucleofector 4D (program B1). Afterwards the cells were diluted into mTeSR-plus media and plated in 90mm dishes coated with Matrigel. 24-hours later, transfection was verified by GFP fluorescence and blasticidin (mg/ml) was added to the media for 48 hours and then removed. Over the following days, individual colonies were picked and re-plated. When sufficient expansion occurred cells were harvest for gDNA that was sequenced to determine *LMNA*-R541C correction (sequencing primers: forward-GTCCCTCTGGGGTGGAAATG, reverse-AGGCCAGCGAGTAAAGTTCC).

*LMNA*-R541C Knock-in cell lines were generated at Synthego Corporation (CA). Electroporation of CRISPR-Cas9 with specific guide RNA and donor oligonucleotide DNA into the PGP1 iPSC line.

Guide RNA: GGAAGUGGCCAUGCGCAAGC.

Donor Sequence:

GACCCTTGGACCTGGTTCCATGTCCCCACCAGGAAGTGGCCATGTGCAAGCTGGTGCGCTCAGTGACTGTGGTTGAGGACGACGAGGATGA.

Clonal cell lines were obtained by limiting dilution of single cell suspensions. Validation of heterozygous introduction of *LMNA*-R541C was determined by Sanger sequencing of genomic DNA PCR products using the following primers: Forward-GGAGAGCTTGACAGTGTCCCC, Reverse-AGGCCAGCGAGTAAAGTTCC.

### Sanger Sequencing

iPSC genomic DNA was isolated using the QIAamp® DNA Mini Kit (Qiagen) and amplified using DREAMTaq PCR MasterMix (ThermoFisher). Resulting sample was run on 1.5% agarose gel at 60v for 1.5-2 hours then purified using Monarch DNA Gel Extraction Kit (New England BioLabs). Samples were Sanger sequenced and correct *LMNA* sequences were confirmed.

### Cardiomyocyte differentiation of iPSC

iPSC lines were differentiated towards ventricular cardiomyocytes (iCMs) using STEMdiff™ Cardiomyocyte Differentiation Kit (STEMCELL Technologies), as recommended by the manufacturer. Briefly, iPSCs were suspended as single cells with Gentle Cell Dissociation reagent (STEMCELL Technologies) and seeded on Matrigel-coated 12-well plates at a density of approximately 90-95% on the day prior to starting differentiation. Cells were then treated with STEMdiff™ Cardiomyocyte Differentiation Medium A supplemented with Matrigel (1:100) for 2 days, STEMdiff™ Cardiomyocyte Differentiation Medium B for 2 days, STEMdiff™ Cardiomyocyte Differentiation Medium C for 4 days, and STEMdiff™ Cardiomyocyte Maintenance Medium, thereafter. 2 weeks from the start of differentiation, purification of cardiomyocytes was achieved by a metabolic-selection method as previously described (Sharma et al., 2015) with glucose starvation (RPMI-glucose + B27) for 5 days before functionality analysis. Fresh media was renewed every other day. After glucose-starvation, iCMs were further cultured in STEMdiff™ Cardiomyocyte Differentiation Medium for a minimum of 30 days from differentiation for enhanced maturity (Karbassi et al., 2020)prior to further analyses. Cardiomyocyte aggregates were dissociated using Cardiomyocyte Dissociation media (STEMCELL Technologies) for downstream experiments.

### RNA extraction for sequencing and qPCR analysis

After maturing cardiomyocytes for at least 30 days RNA extraction was done using RNeasy Mini Kit and QIAshredder™ (Qiagen) for lysis according to manufacturer’s instructions. Concentration and purity of RNA samples was measured using a Nanodrop before downstream sequencing or use in qPCR.

For each experiment, 3 replicates for each treatment were sent to Novagene Co. Ltd for bulk mRNA sequencing. Total RNA (1 μg) from each sample was utilized to generate sequencing libraries using the NEBNext UltraTM RNA Library Prep Kit for Illumina (NEB, USA). mRNA was enriched from total RNA by poly-T oligo-attached magnetic beads. Double-stranded cDNA synthesis was accomplished by using M-MuLV Reverse Transcriptase, random hexamer primers, Polymerase I and RNase H. AMPure XP technology was used to select cDNA fragments with a preferred length of 150–200 bp (Beckman Coulter, Beverly, USA). Samples were then incubated with USER Enzyme (NEB, USA) for 15 minutes at 37 °C, followed by 5 minutes at 95 °C. Next, PCR was carried out using Phusion High-Fidelity DNA Polymerase, Universal PCR primers, and Index (X) Primer. Lastly, PCR products were purified using the AMPure XP system and the library quality was determined using the Agilent Bioanalyzer 2100 system. Clustering the index-coded samples was performed on a cBot Cluster Generation System using PE Cluster Kit cBot-HS (PE-401-3001, Illumina, CA, USA). After cluster generation, the library preparations were sequenced on an Illumina platform and paired-end reads were produced.

For qPCR, cDNA was made using 150-200ng of RNA using the SuperScript™ IV VILO™ MasterMix kit (Invitrogen) according to manufacturer’s instructions. qPCR was done with 15ng cDNA per reaction with the TaqMan Fast Advanced MasterMix (ThermoFisher) and TaqMan gene expression probes (ThermoFisher). Expression was normalized to GAPDH and compared to wild-type (PGP-1 or unrelated control cells) using the ΔΔCT method.

### Computational analyses

The raw fastq data was quantified using Salmon (v1.10.3 (Patro et al., 2017)) with a decoy aware index using the hg38 human genome in Ensembl (v97 (Harrison et al., 2024)).

### Statistical analyses

All statistical analyses were performed in R (v4.4.1).

### Differential expression

Initially, each experiment was analyzed independently, comparing *LMNA*^WT^ to *LMNA*^R541C^ (Fig. 1A). For gene level analyses, transcript expression was summarized to the gene level using the tximport (Soneson et al., 2015) summarizetogene method using the most abundant isoform to define gene ranges. DESeq2 (Love et al., 2014) was used to perform gene-level differential expression analyses, and the vst function was used to scale the data for use in downstream visualization. P values were adjusted using the Benjamini-Hochberg correction (Benjamini C Hochberg, 1995). Gene lists for downstream analyses were constructed using all genes with an absolute log2 fold-change > 1 and adjusted p-value < 0.05.

**Figure 1.**
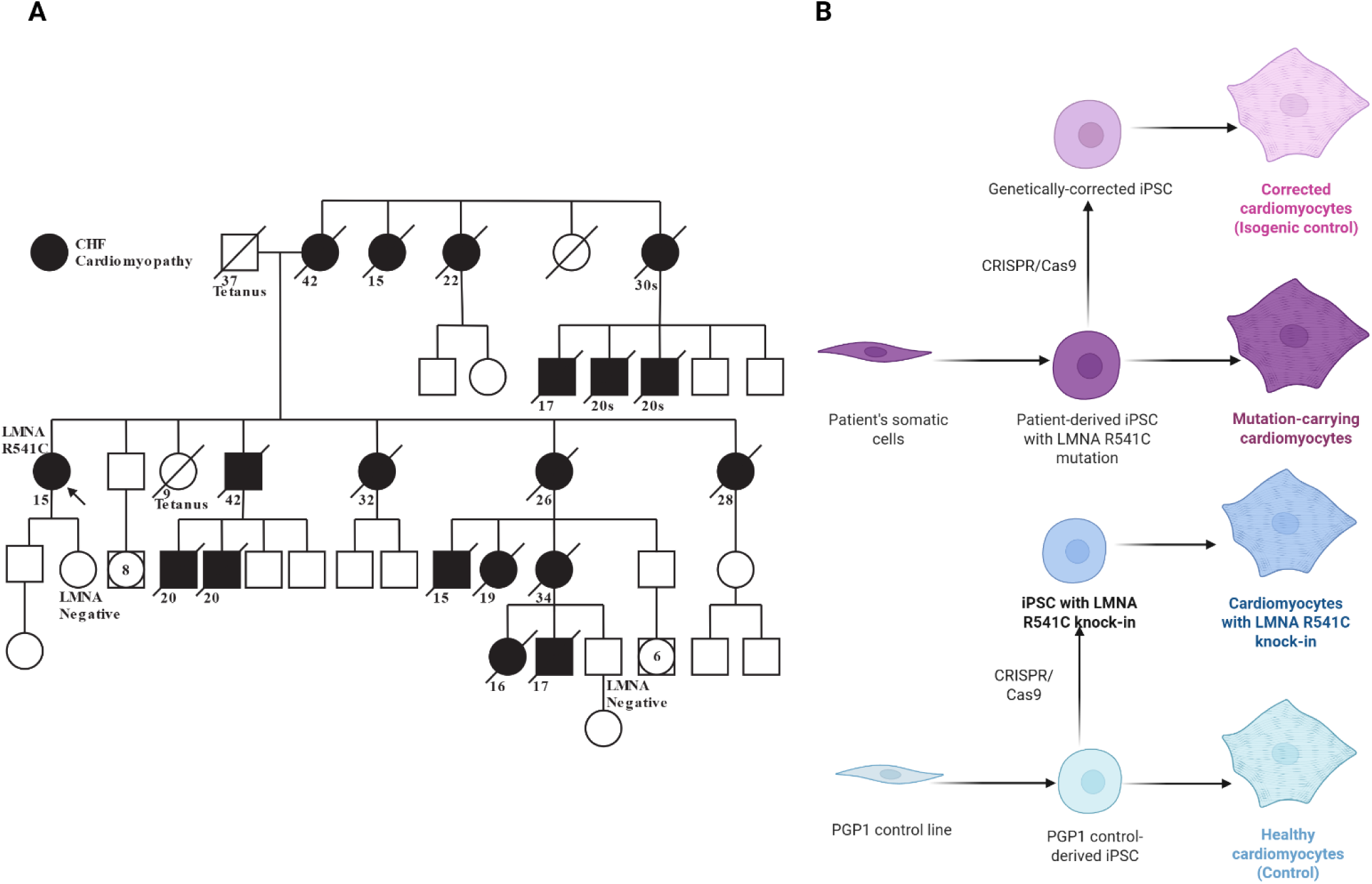
Experimental design and family history of highly penetrant and severe phenotype of *LMNA*^R541C^. Multi-generational genogram of family demonstrating prevalence of cardiomyopathy with early lethality. Numbers indicate onset of heart failure or the age of death designated by the diagonal lines. B) Schematic of experimental design for Crispr-correction and Crispr knock-in of *LMNA*-R541C mutation.

### Pathway enrichment

Gene pathways were collected from the Gene Ontology database (GO:BP (Ashburner et al., 2000)), and Molecular Signatures Database (MSigDB) (Hallmark and C2 (Subramanian et al., 2005)). Enriched pathways were then identified using ranked gene lists according to log2 fold change values with the Gene Set Enrichment Analysis (GSEA) function from the clusterProfiler (Yu et al., 2012) R library. Enrichment results were also visualized with the GeneTonic library (Yu et al., 2012).

Gene Set Variation Analysis (GSVA (Hanzelmann et al., 2013)) was used to calculate per-sample pathway enrichment using Gene Ontology and MSIG as source databases. The gsva and ssgsea single-sample methods were used to calculate enrichment. For downstream plotting and reporting, we considered pathways that were identified as enriched by both methods. After pathway enrichment values were calculated, limma (Kramer et al., 2014) was used to identify differential pathway enrichment. The regression model used for limma was “∼ experiment + mutation” to account for differences between experiments. We also used the gene-set size as a trend variable when calculating significance with the eBayes function.

Pathway enrichment was also conducted with Ingenuity Pathway Analysis to identify and visualize cardiovascular disease-specific pathways and networks (Ritchie et al., 2015).

### Drug response prediction using LINCs database

The drug response LINCS L1000 database (Subramanian et al., 2017) was used to identify potential drug targets for *LMNA* R541C (Kwee et al., 2022). The R package metaLINCS was used to test for drugs and pathways with similar gene perturbations to the LMNA experiments (Kwee et al., 2022).

## Results

### Identification of *LMNA* R541C in a familial cardiomyopathy cohort

A pathogenic *LMNA* variant (c.1621C>T, p.R541C, heterozygous) was observed in a middle-aged woman with severe congestive heart failure with reduced ejection fraction requiring heart transplantation. She had experienced symptoms of heart failure since her adolescence. The family history was remarkable strong for a Mendelian inheritance pattern with high degree of penetrance and early onset of severe cardiomyopathy with high lethality (Genogram in Figure 1A). Peripheral blood mononuclear cells from this patient were reprogramed into iPSCs (Fig. 1B) as previously reported (Yang, Argenziano, et al., 2021; Yang, Burgos Angulo, et al., 2021).

### Individual differential expression and pathway analysis of experiments

We first analyzed each experiment independently. Using DESeq2 there were 3699 and 330 genes significantly differentially expressed and with a minimum absolute log2 fold change >=1 in the CRISPR-corrected, knock-in, and PGP-R541C experiments, respectively. Volcano plots and heatmaps for the DEGs are shown in Figure 2. There was considerable variation between experiments in the individually DE genes(N=68 and N=55 for the shared up- and down-regulated genes, Supplementary Table 1).

**Figure 2.**
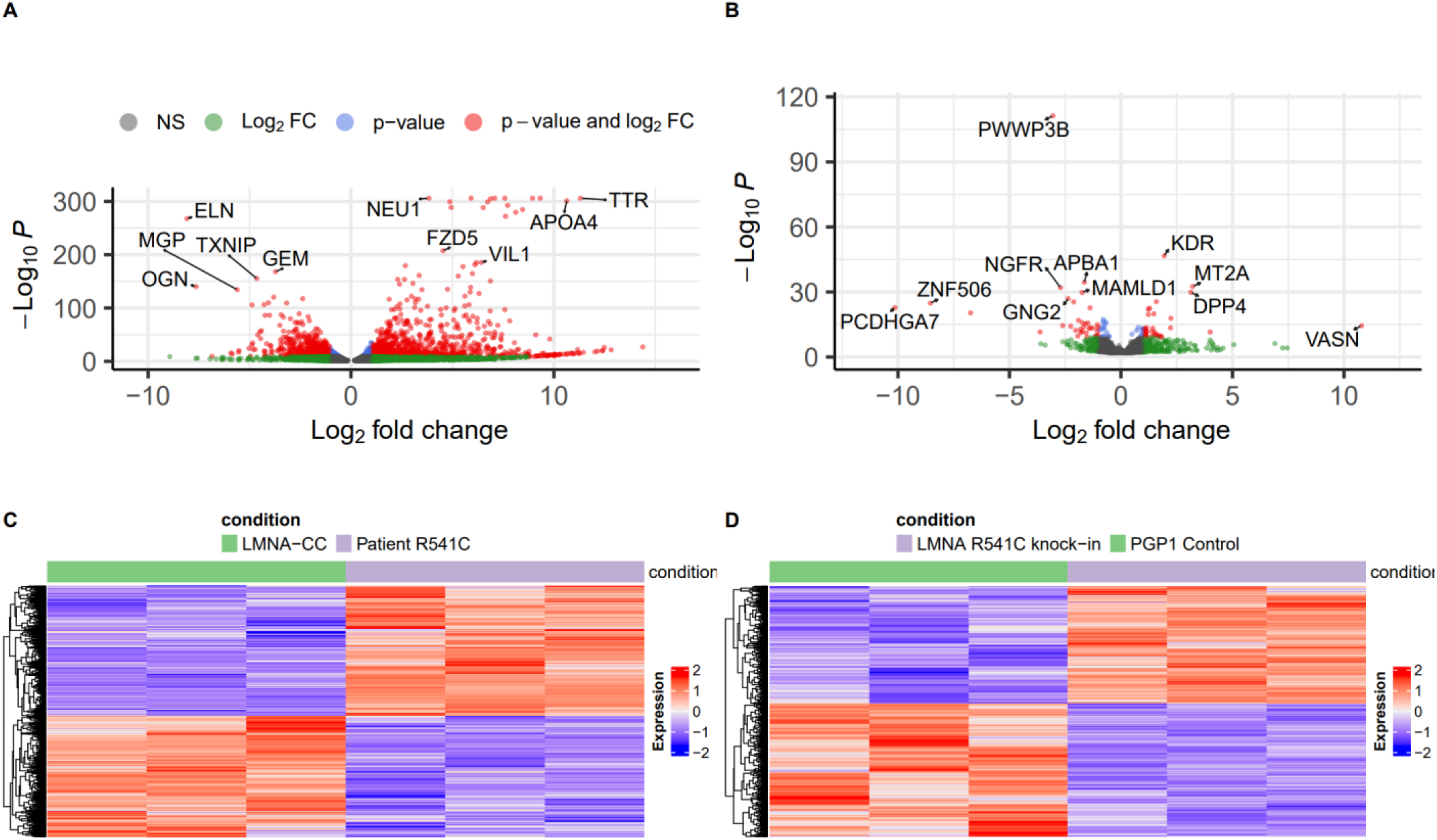
Differential gene expression for heterozygous *LMNA*-R541C (*LMNA*^R541C/WT^ vs *LMNA*^WT/WT^). *LMNA*-R541C CRISPR rescue and knock-in experiments indicate gene rescue effects and gene dysregulation in iPSC cardiomyocytes. A) Volcano plot of differentially expressed genes in *LMNA* CRISPR-correction experiment. B) Volcano plot of differentially expressed genes in *LMNA* knock-in experiment. C) Heatmap for *LMNA-*CRISPR-corrected experiment. D) Heatmap for *LMNA* knock-in experiment.

### Variation in differentially expressed genes between experiments

Our initial comparison of the different experiments identified genes that were up- or down-regulated in both CRISPR-corrected and knock-in analyses. For both up- and down-regulated genes, the number of common genes was smaller than the number unique to either experiment. Batch effects between experiments were accounted for by explicitly included experiment as a variable in models.

### GSVA analysis reveals common enriched pathways across experiments

Due to the observed differences between the experiments, we used Gene Set Variation Analysis (GSVA) to calculate pathway enrichment on a per-sample basis. Using a model to partition variance explained by experiment as well as by mutation, we identified pathway changes specifically due to the *LMNA-*R541C compared to *LMNA* WT. The primary analysis considered the CRISPR-correction and knock-in experiments together.

For the gene ontology Biological Process pathways, there were N=673 significant differentially expressed pathways between *LMNA-*R541C and *LMNA* WT. The most significant terms relate to protein translation, the ribosome, and axon guidance (Figure 3A). We also calculated enrichment among the MSigDB Hallmark pathways. Among these terms, there were N=14 significant pathways. The primary enriched Hallmark pathways relate to *MYC* regulation, fatty acid metabolism, and adipogenesis (Figure 3B).

**Figure 3.**
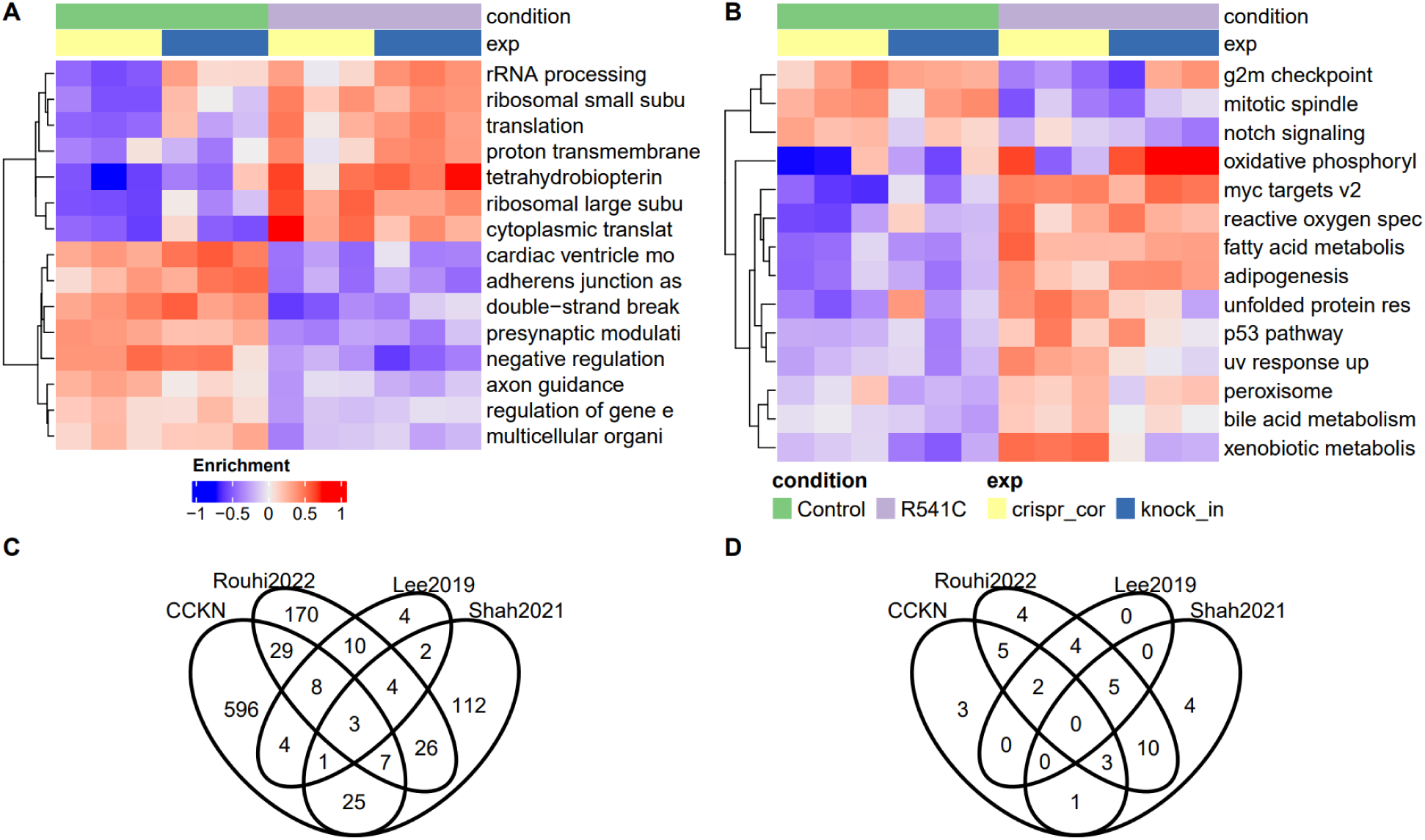
*LMNA* R541C-associated pathways dysregulation. Pathways dysregulated in *LMNA-*R541C are also dysregulated in prior studies with alternate *LMNA* mutations. A) Heatmap of enriched GO:BP pathways using GSVA. B) Heatmap of enriched MSigDB Hallmark pathways using GSVA. C) Overlap of GO:BP pathways and published *LMNA* RNA-seq D) Overlap of MSigDB Hallmark pathways and published *LMNA* studies.

### Differentially regulated genes shared between *LMNA* R541C, *LMNA* K117fs, and *LMNA*-null experiments

We reanalyzed data from three prior *LMNA* studies, two in human iPSC CM cells and one in mice (Lee et al., 2019; Rouhi et al., 2022; Shah et al., 2021). For this comparison, we considered any gene significant after correcting p-values for multiple testing (false-discovery rate). The most significant genes overlapping multiple datasets are shown in Table 1. The entire list of genes overlapping at least one experiment is listed in Supplementary table 2. The most significant genes that were also significant in the (Shah et al., 2021) study are listed in table 2.

**Table 1.**
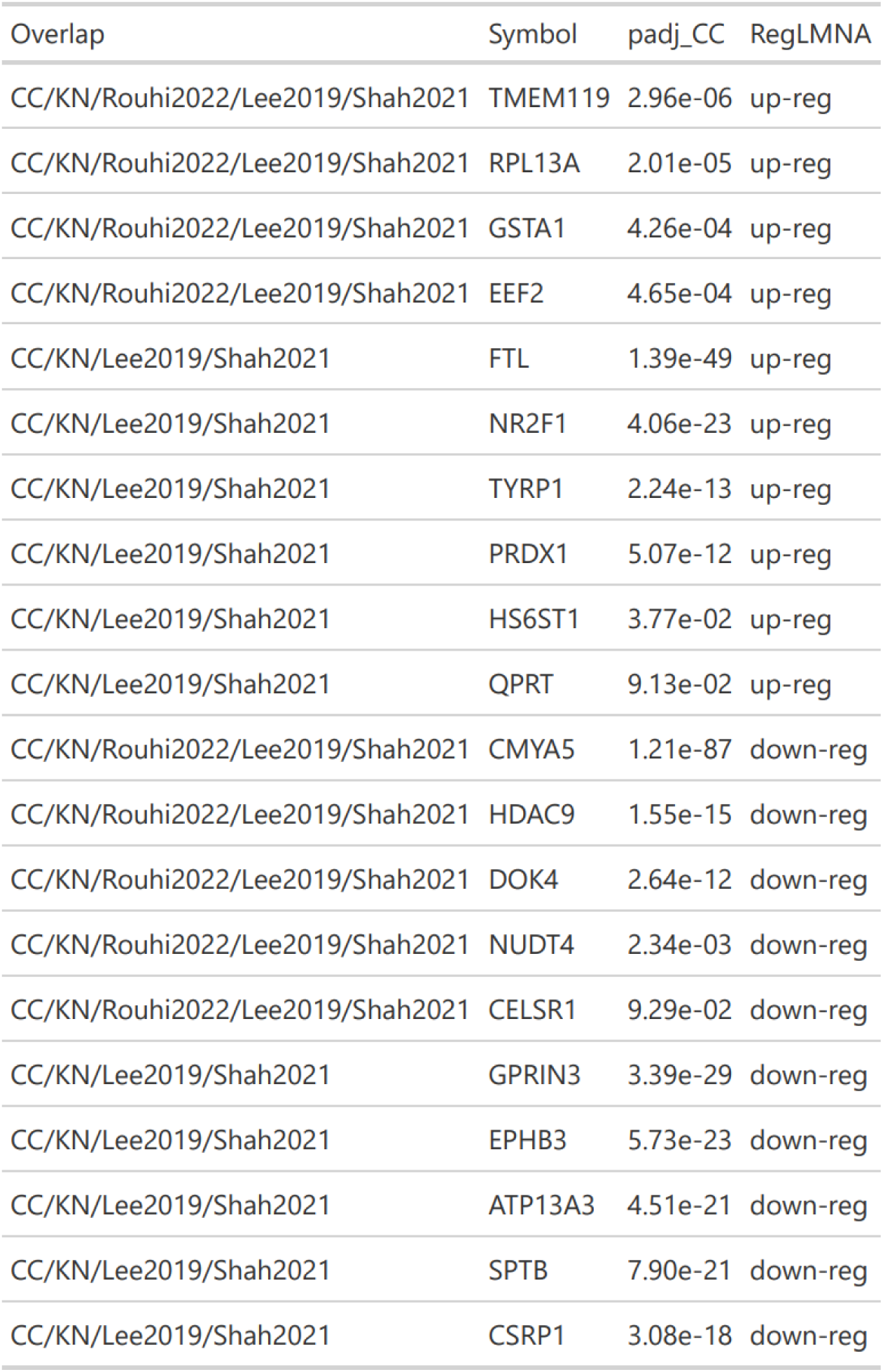
Significant DEG shared by CRISPR/knock-in experiments and prior studies (mouse model and human IPSC model). The 10 most significant up/down regulated genes are shown for each set of study overlaps.

**Table 2.** Substantial overlap of top DEGs with Lee et al. 2019 IPSC study involving *LMNA* frameshift mutation.

| symbol | expr_avg | log2fc_avg | padj_avg | cckn_expr |
| --- | --- | --- | --- | --- |
| CLU | 5450.95848 | 1.9482983 | 5.472058e-18 | upreg |
| FTL | 16159.66447 | 1.2896816 | 5.209274e-11 | upreg |
| PRDX1 | 4317.15386 | 0.7252314 | 3.204730e-09 | upreg |
| HMGA1 | 1752.66058 | 1.2182626 | 9.386676e-07 | upreg |
| HMOX1 | 463.91535 | 3.5502902 | 1.246344e-06 | upreg |
| NDRG1 | 2006.00937 | 1.6374298 | 3.565954e-06 | upreg |
| PLIN2 | 918.77224 | 1.4463031 | 5.954716e-06 | upreg |
| TYRP1 | 74.82981 | 2.3524169 | 1.165538e-05 | upreg |
| MUC13 | 49.42145 | 5.6887751 | 2.201149e-05 | upreg |
| TMEM119 | 157.43427 | 1.3004572 | 4.769108e-05 | upreg |
| MAMLD1 | 510.90961 | -2.1702109 | 2.049561e-14 | downreg |
| ELOVL5 | 2880.04279 | -1.0149404 | 3.069926e-14 | downreg |
| SLITRK4 | 421.19062 | -1.6873854 | 7.511092e-14 | downreg |
| COL8A1 | 3359.87468 | -2.8053819 | 1.463145e-13 | downreg |
| HEYL | 357.64384 | -2.4868847 | 1.664624e-12 | downreg |
| AFAP1 | 3962.79354 | -1.1363198 | 4.182468e-12 | downreg |
| JAG1 | 3494.00271 | -1.1467252 | 1.541957e-11 | downreg |
| HIP1 | 2987.31677 | -0.8054997 | 9.651349e-10 | downreg |
| MICAL2 | 14107.89210 | -0.9707714 | 2.752884e-09 | downreg |
| BMPR2 | 6412.50399 | -0.6924999 | 4.592155e-09 | downreg |

### Shared pathways between *LMNA* R541C, *LMNA* K117FS and *LMNA* -/- involved in proliferation, metabolism, and development

We identified shared GO:BP and MSigDB Hallmark pathways after performing GSEA analysis on the different *LMNA* experiments (Figure 3C-D, Table 3, Supplemental Table 3). There were 77 shared biological processes between the *LMNA* R541C and prior *LMNA* experiments, but most (29/77) were only enriched in data from (Rouhi et al., 2022) or (Shah et al., 2021) (25/77); 3 pathways were shared across all 4 experiments. Most (10/11) of the significant Hallmark pathways identified in the CRISPR-corrected and knock-in experiments were also identified in (Rouhi et al., 2022). Overall, these pathways relate to 3 broad categories defined in (Subramanian et al., 2005), specifically proliferation, metabolism, and development.

**Table 3.** Pathways conserved between CRISPR-corrected/knock-in experiments (CCKN), mouse models (Rouhi2022), and human IPSC (Lee2019, Shah2021) studies.

| Overlap | Pathway | Process Category |
| --- | --- | --- |
| CCKN/Rouhi2022/Shah2021 | fatty acid metabolism | metabolic |
| CCKN/Rouhi2022/Shah2021 | mitotic spindle | proliferation |
| CCKN/Rouhi2022/Shah2021 | bile acid metabolism | metabolic |
| CCKN/Rouhi2022/Lee2019 | oxidative phosphorylation | metabolic |
| CCKN/Rouhi2022/Lee2019 | g2m checkpoint | proliferation |
| CCKN/Shah2021 | xenobiotic metabolism | metabolic |
| CCKN/Rouhi2022 | adipogenesis | development |
| CCKN/Rouhi2022 | uv response up | DNA damage |
| CCKN/Rouhi2022 | p53 pathway | proliferation |
| CCKN/Rouhi2022 | peroxisome | cellular component |
| CCKN/Rouhi2022 | unfolded protein response | pathway |
| CCKN | myc targets v2 | proliferation |
| CCKN | reactive oxygen species_pathway | pathway |
| CCKN | notch signaling | signaling |

### Cardiovascular-specific disorders are enriched in consistently dysregulated *LMNA* R541C genes

We used the previously identified list of consistently dysregulated genes to conduct pathway enrichment using Ingenuity Pathway Analysis. The most enriched general pathway was preeclampsia signaling, followed by several pathways associated with non-cardiovascular organs (Figure 4A). In addition, a collection of cardiovascular-specific disease pathways was also enriched (Figure 4B). The cardiovascular pathways were predominantly related to hypertension and abnormal heart morphology. A graphical network summary highlighted a dysregulated pathway involving TGFBR1, PPARG, SMARCA4, TNF, and others (Figure 4C). Together, these results show further support for the general biological pathways identified, as well as providing cardiovascular-specific disease pathways.

**Figure 4.**
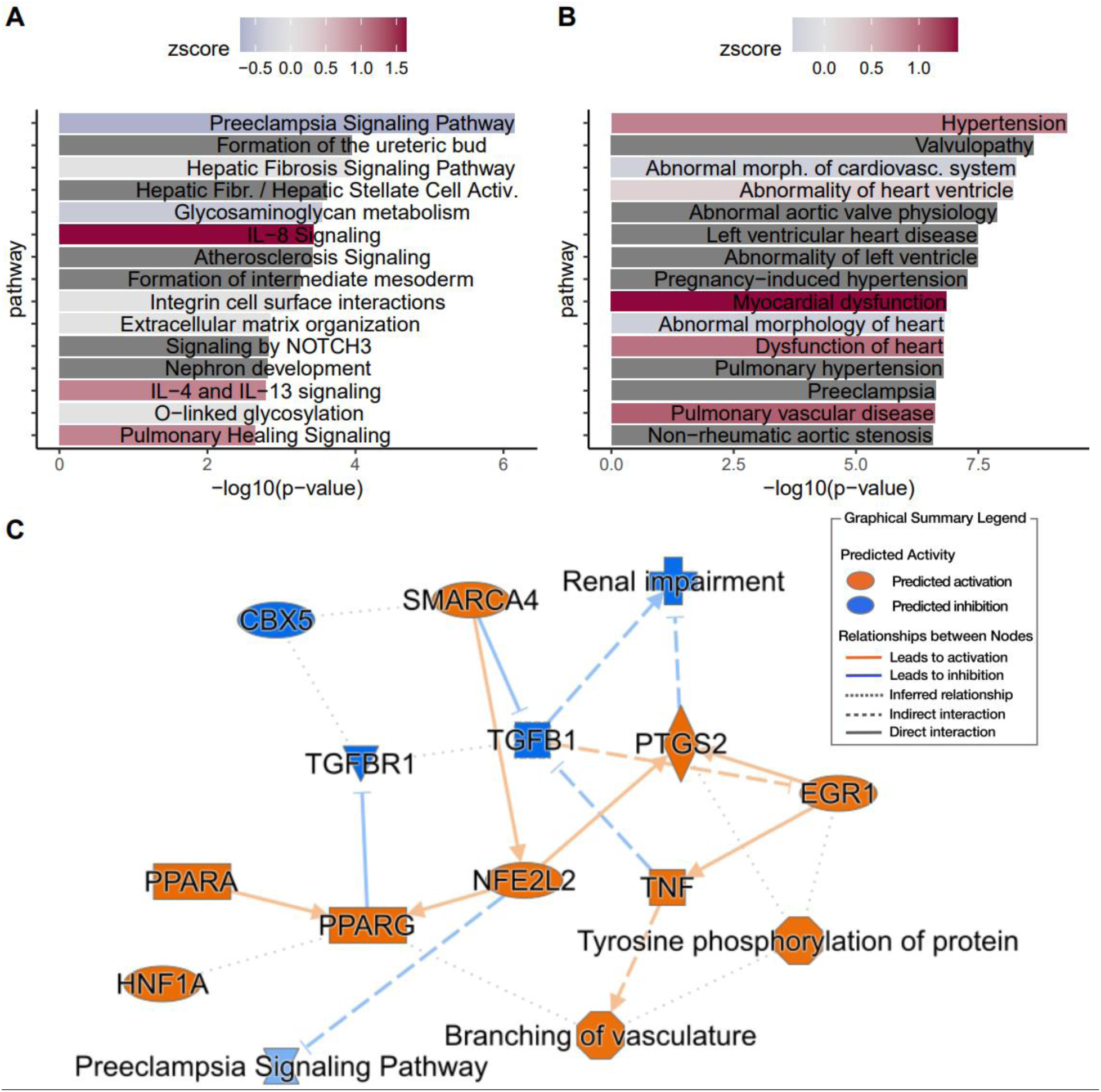
Ingenuity Pathway Analysis of differentially expressed genes associated with *LMNA-*R541C. Bar colors indicate normalized z-score enrichment (positive= overexpression in LMNA-R541C, negative=underexpressed in LMNA-R541C, dark grey= directionality could not be ascertained by IPA). A) IPA canonical pathway enrichment. B) Enrichment of cardiovascular disease-specific pathways. C) Gene-pathway network summary of most significant associations.

### Drug-response prediction identifies heterogenous response to drugs among experiments

Finally, we used the calculated fold-change of all genes to identify potential drug targets using metaLINCS (Kwee et al., 2022). There was substantial variation across the experiments among the most-enriched pathways and genes across the LINCs database (Figures 5A-D), with only the estrogen receptor agonist pathway among the most enriched pathways for both experiments. However, examining the predicted drug response for the top-ranked drugs reveals a subset with consistent drug activity mapping (Figure 5E). The top predicted compensatory drug targets included cardiac glycoside-related compounds and mTOR kinase inhibitors.

**Figure 5.**
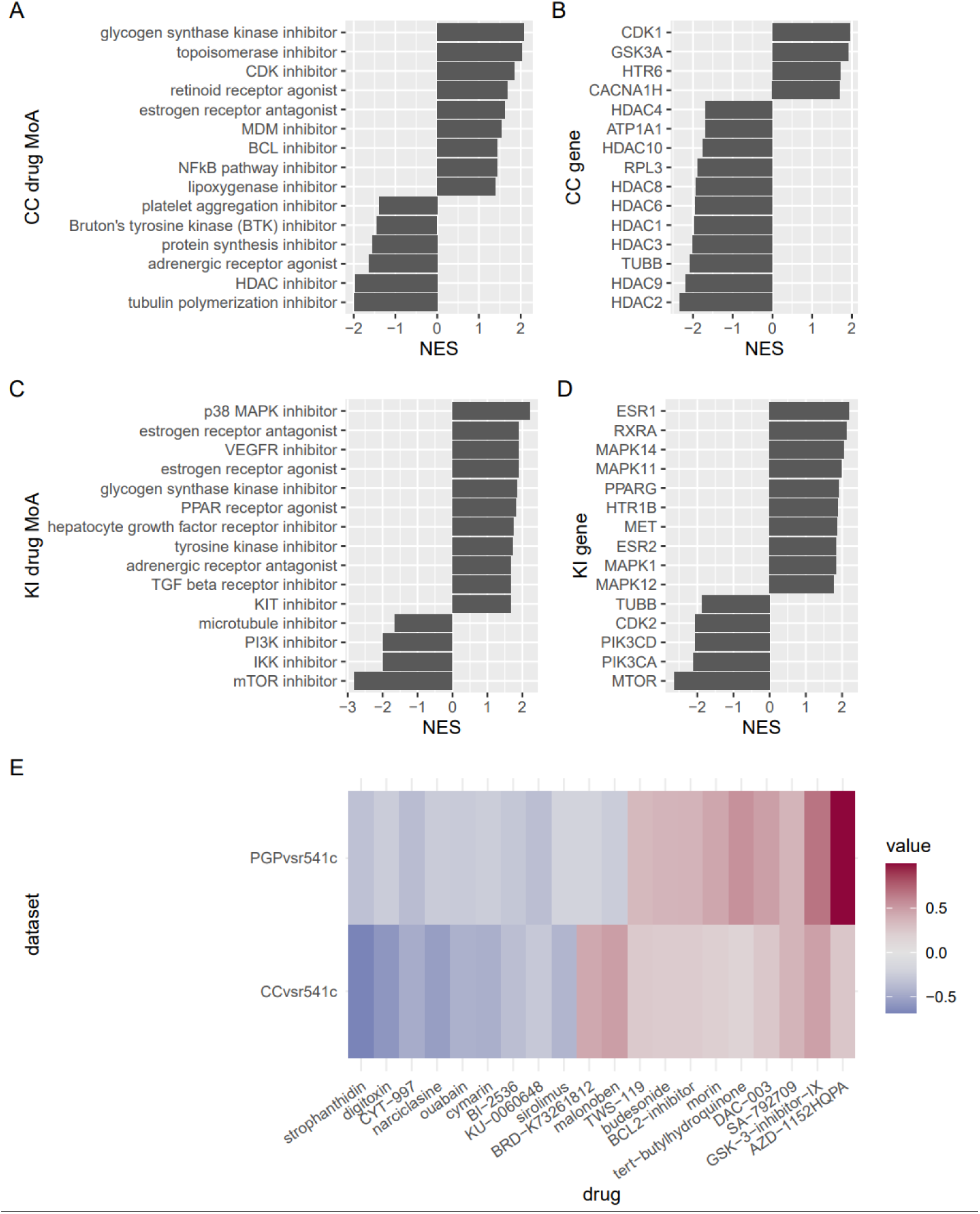
Meta-level analysis of LINCS drug signatures. metaLINCS database screen identified drugs classes with shared response from *LMNA-*R541C differential gene expression. Negative values are associated with pathways, genes and drugs acting in potentially compensatory fashion to LMNA mutation. A) drug mechanism of action (MoA) enrichment in CRISPR-corrected experiment. B) Gene enrichment in CRISPR-corrected data. C) drug mechanism of action (MoA) enrichment in knock-in experiment. D) Gene enrichment in knock-in experiment. E) Drug activity map identify drugs with similar activity across experiments. Values (color) are the relative activation across test profiles in the LINCS database. Negative values are anti-correlated with *LMNA*-R541C changes and potentially compensatory, while positive values are positively correlated with *LMNA*-R541C changes and potentially exacerbating.

## Discussion

In this study we used complementary CRISPR-based approaches to model the *LMNA*-R541C pathogenic variant in human iPSC-derived cardiomyocytes and to isolate its genotype-specific transcriptional effects. We constructed iPSC cell lines with isogenic controls where *LMNA-*R541C was either CRISPR-corrected to wild-type in patient-derived *LMNA*^R541C/WT^ cells or introduced as a heterozygous knock-in with a control iPSC line. RNAseq analysis of these iPSCs that had been differentiated into cardiomyocytes showed 123 genes that were both significantly and consistently dysregulated in the same manner across experiments. Pathway analysis identified 77 shared pathways from Gene Ontology biological processes, and 14 shared pathways from the MSigDB Hallmark database. Together, these results describe the early molecular and biological consequences that may initiate a subsequent severe and aggressive cardiomyopathy arising from a single amino acid substitution in *LMNA*.

There were numerous genes consistently altered in the CRISPR-correction and knock-in experiments. The genes with the highest consistent fold-change values were *GSTA1, MUC13, CEACAMc, CDH17*, and *VIL1. GSTA1* is an enzyme that transfers glutathione to compounds which can detoxification, lipid metabolism, and oxidative stress responses; recent Mendelian randomization studies indicated that this gene is associated with cardiovascular function (Jiang et al., 2024; Selber-Hnatiw et al., 2026). Several are associated with oxidative stress responses, epithelial signaling pathways, and cellular detoxification processes. *MUC13* and *CEACAMc* are cell-surface glycoproteins which are usually lowly expressed in cardiovascular tissues (Williams et al., 2001; Zhao et al., 2024) consistent with the hypothesis of pathogenic *LMNA* variants resulting in misexpression of genes not normally expressed in a given tissue (Shah et al., 2021).

GSEA pathway analysis revealed multiple highly significant pathways involved in metabolism, cell proliferation, and development. Notably, 11 of the 14 pathways enriched from the current experiments (such as fatty acid metabolism and mitotic spindle) were also enriched in previously published *LMNA* RNA-seq based analysis (Lee et al., 2019; Rouhi et al., 2022; Shah et al., 2021). The enriched Hallmark pathways in the current experiments were among the most significantly altered pathways (with *MYC* targets v2 being the most significantly changed across all pathways, FDR= 4.9e-5, log2FC = 0.56). Additional distinct pathways were reactive oxygen species (ROS), xenobiotic metabolism, and notch signaling at ranks 4, 5, and 8 out of 14 total enriched pathways. *MYC* regulation, ROS, and notch signaling have all been previously observed in cardiovascular development and disease (Aquila et al., 2013; Moris et al., 2017; Napoli et al., 2002). We also calculated pathway enrichment using Ingenuity Pathway Analysis (IPA, Qiagen). IPA suggested that the most enriched general term was preeclampsia signaling. Other notable pathways were glycosamingoglycan (extracellular matrix), IL-8 (inflammation), and atherosclerosis. The consistent identification of G2/M checkpoint dysregulation across multiple *LMNA* models suggests that nuclear structural defects may perturb cell cycle control and DNA damage responses, processes increasingly recognized as contributors to cardiomyocyte dysfunction (Wallace et al., 2023; Zhao et al., 2026).

One of the 3 external datasets we compared was (Shah et al., 2021), which was also an iPSC comparison of *LMNA*-R541C. Of the consistently differentially expressed genes in the CRISPR-correction and CRISPR knock-in experiments, more than half (329/646) were up- or down-regulated in a similar expression in (Shah et al., 2021). Interestingly, although there was some overlap in pathways between our experiments and (Shah et al., 2021), 89% (596/673) of the enriched GO:BP pathways were not significantly changed in other experiments. Further study of *LMNA*-R541C is warranted to elucidate the genes and pathways that are consistently affected across experiments, which may indicate targets for intervention.

Dysregulation of oxidative phosphorylation pathways may reflect mitochondrial dysfunction secondary to altered nuclear-mitochondrial signaling. Cardiomyocytes rely heavily on mitochondrial ATP production, and impairment of oxidative phosphorylation has been reported in multiple genetic cardiomyopathies (Bhullar & Dhalla, 2023). *LMNA*-associated nuclear defects may therefore indirectly influence mitochondrial metabolism through altered transcriptional programs or mechano-transduction pathways.

Several potential drug targets were informatically identified using the LINCS drug perturbation database. Candidate molecules for these targets include strophanthidin, digitoxin, CYT-997 (lexibulin), narciclasine, ouabain and cymarin. Several of these (strophanthidin, digitoxin, ouabain) are related compounds known as cardiac glycosides that have been used in symptomatic congestive heart failure and arrhythmias but are limited by narrow therapeutic index. Cardiac glycoside therapy is controversial; a recent meta-analysis of 6 glycoside studies indicated glycoside administration reduced heart-failure hospitalizations, but not all-cause mortality (Wang et al., 2026). Digitoxin in a recent double-blind trial, was associated with lower risks of death in patients with heart failure and reduced ejection fraction (Bavendiek et al., 2025). Whether digitoxin possesses other unrecognized targets beyond those established for cardiac glycosides (Arispe et al., 2008) is yet to be determined. Aside from the established cardiac glycosides, narciclasine has anti-inflammatory properties and led to improved cardiac function in mouse models of sepsis-induced myocardial dysfunction (Tang et al., 2025).

A limitation of this study includes solely analyzing cardiomyocyte iPSCs. Previous studies identified variation between cardiomyocytes and cardiofibroblasts with *LMNA*^R541C^ (Yang, Argenziano, et al., 2021), so the underlying molecular pathways may also differ in cell-specific ways. Furthermore, the potential drug targets were datamined using the LINCS database and have not been directly tested in the *LMNA* iPSC models yet.

In summary, our results identify consistent transcriptional and pathway-level alterations associated with the *LMNA*-R541C mutation in human cardiomyocytes. These findings implicate coordinated dysregulation of cell-cycle control, mitochondrial metabolism, and stress signaling pathways as early molecular consequences of *LMNA* dysfunction. Future studies integrating additional *LMNA* variants, multiple cardiac cell types, and functional perturbation experiments will be necessary to further define the mechanisms linking nuclear lamina defects to cardiomyopathy progression and to identify precision therapeutic opportunities for *LMNA*-associated heart disease.

## Acknowledgements

Figure 1 created in BioRender. Keller, T. (2026) https://BioRender.com/jggfkb5 Grant support: NIH/NHLBI 1R56 HL156146 to TVM.

## Data availability

The sequence data for the CRISPR-correction and knock-in experiments are located at GSE334682.

## Code availability

All R code for data analysis and figure generation are available at https://github.com/thomas-keller/lmna_r541c_paper

